# Aneuvis: Web-based exploration of numerical chromosomal variation in single cells

**DOI:** 10.1101/459735

**Authors:** Daniel G Piqué, Grasiella A Andriani, Elaine Maggi, Samuel E Zimmerman, John M Greally, Cristina Montagna, Jessica C. Mar

## Abstract

**Motivation:** Aberrations in chromosomal copy number are one of the most common molecular features observed in cancer. Quantifying the degree of numerical chromosomal variation in single cells across a population of cells is of interest to researchers studying whole chromosomal instability (W-CIN). W-CIN, a state of high numerical chromosomal variation, contributes to treatment resistance in cancer.

**Results:** Here, we introduce aneuvis, a web application that allows users to determine whether numerical chromosomal variation exists between experimental treatment groups. The web interface allows users to upload molecular cytogenetic or processed whole-genome sequencing data in a cell-by-chromosome matrix format and automatically generates visualizations and summary statistics that reflect the degree of numeric chromosomal variability. Aneuvis is the first user-friendly web application to help researchers identify the genetic and environmental perturbations that promote numerical chromosomal variation.

**Availability and Implementation:** Aneuvis is freely available as a web application at https://dpique.shinyapps.io/aneuvis/. Website implemented using Shiny version 1.0.5 with all major browsers supported. All source code for the application is available at https://github.com/dpique/aneuvis.

## 1. Introduction

Alterations in chromosome number are a hallmark of cancers (Taylor *et al.*, 2018). Within a population of single cells, increased numerical chromosomal variation may reflect underlying whole chromosomal instability (W-CIN) (Geigl *et al.*, 2008), which promotes chemotherapy resistance (Sansregret *et al.*, 2018). The process of identifying numerical chromosomal variation in single cells can be divided into two steps. The first step is to quantify the number of chromosomes per cell. Multiple experimental techniques and computational tools exist for completing this step, and the final output is often a spreadsheet or text file that contains chromosomal copy number information for all nuclei analyzed (Bakker *et al.*, 2015). The second step is to quantify the degree of numerical chromosomal variation. Existing approaches, such as AneuFinder, estimate the degree of numerical aneuploidy from low coverage single cell whole genome sequencing (scL-WGS) data but require knowledge of the R programming language (Bakker *et al.*, 2016). In addition, existing approaches do not allow specification and comparison of experimental treatment groups. Furthermore, no freely-available software exists for the processing of chromosomal count data derived from locus specific fluorescent *in situ* hybridization (FISH) or spectral karyotyping (SKY).

Here, we introduce aneuvis, a user-friendly web application for visualizing and summarizing numerical chromosomal variability in populations of single cells belonging to experimentally-defined treatment groups. Aneuvis operates downstream of existing experimental and computational approaches that generate a matrix containing the estimated chromosomal copy number per cell. Users upload both a copy number matrix along with a key that links individual cells with experimental treatment groups. Aneuvis is the first freely available, user-friendly application to automatically calculate metrics and generate graphics that reflect numerical chromosomal variation between experimental treatment groups. Aneuvis is available to use as a standalone web application at https://dpique.shinyapps.io/aneuvis/.

## 2 Materials and Methods

### Input

Aneuvis accepts single cell chromosomal copy number data from different treatment conditions **(Figure 1A).** Three types of single-cell chromosomal data can be uploaded into aneuvis **(Supplementary Figure 1** shows the data upload interface). Fluorescence *in situ* hybridization (FISH) or spectral karyotyping (SKY) data can be uploaded as a Microsoft Excel file where each column is a chromosome and each row is a separate biological cell (or *vice versa*, see **Figure 1B),** and single cell whole genome sequencing (sc-WGS) data at any coverage can be uploaded as a bed file that contains integer estimates of the copy number across the genome. An additional file that links each cell to a treatment group is required and is described within the application.

**Figure 1.**
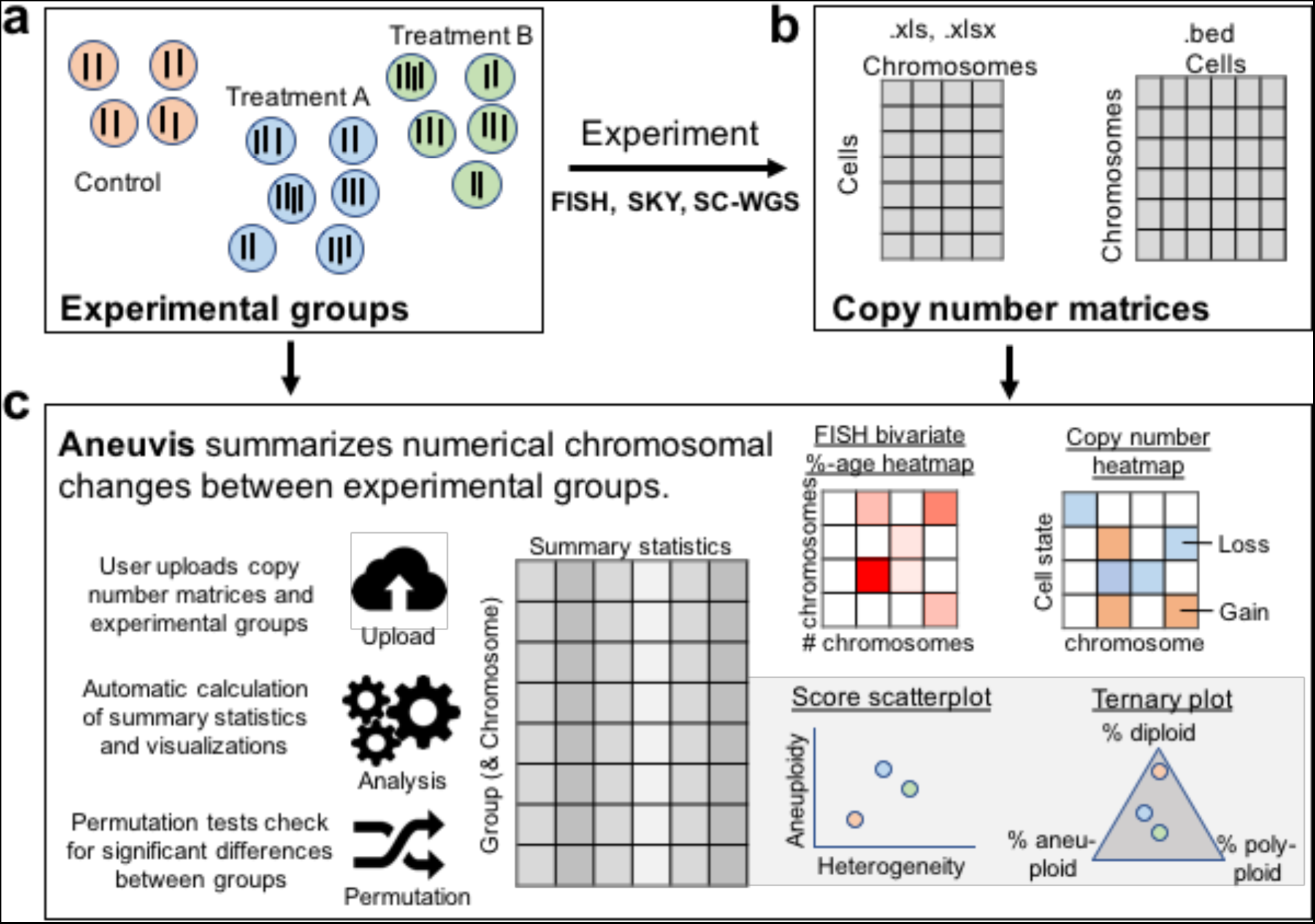
Overview of aneuvis workflow for analyzing numerical chromosomal variation. A) An experiment begins with the quantification of number of chromosomes per cell using either FISH, SKY, or sc-WGS. B) Next, the number of chromosomes per cell within each treatment group is stored as a cell x chromosome matrix, where the entries indicate the number of inferred copies of a chromosome in a cell. C) Aneuvis incorporates information from the experimental design as well as from chromosomal copy number matrices to determine whether differences exist between treatment groups. A table of descriptive statistics summarized by group and by chromosome is automatically generated and available for download. Visual representations of the relationship in aneuploidy between different groups are also automatically generated. A permutation-based approach enables the user to conclude whether there is a statistically significant difference in the ploidy characteristics between treatment groups.

### Output

The output from aneuvis is organized into three sections – table summary, visualization, and hypothesis testing – that are generated automatically from the uploaded data. The table summary includes statistics that summarize the degree of numerical chromosomal aneuploidy and heterogeneity in a population of single cells (see **Supplementary Methods**). The visualization tab includes scatterplots and heat maps derived from the statistics available in the table summary. These visualizations enable comparisons between experimental groups and across experimental approaches. The hypothesis testing section allows users to ask whether differences in summary statistics exist between groups using permutation testing. A heatmap visualization of the permuted output is automatically generated.

### Licensing and availability

Aneuvis was created using Shiny version 1.0.5 (R version 3.4.3) and is available under a GPLv3 license at https://dpique.shinyapps.io/aneuvis/. Source code is available on Github (https://github.com/dpique/aneuvis), and a video tutorial is available within the application.

## 3 Usage scenario

Replicative senescence of mammalian cells is associated with W-CIN *in vitro*, as assessed by four-color interphase FISH (Andriani *et al.*, 2016). To demonstrate the utility of aneuvis, the copy number status of high-passage IMR90 fibroblasts (i.e. replicative senescent fibroblasts) were compared with low-passage fibroblasts (i.e. young fibroblasts) using two techniques: four-color interphase FISH and scL-WGS (Supplementary Methods). Permutation testing from aneuvis revealed that senescent fibroblasts were significantly different from young fibroblasts in terms of the aneuploidy and heterogeneity scores (Bakker *et al.*, 2016) derived from FISH (500 permutations, p = 0.002) but not using sc-WGS (500 permutations, p > 0.05). These results were supported by the FISH bivariate heatmaps, which show increased numerical chromosomal variation in senescent fibroblasts relative to young fibroblasts (Supplementary Figure 3). Inconsistencies between FISH and scL-WGS in measuring aneuploidy are recognized, and likely due to a differential sensitivity of these techniques (FISH is prone to detection of false positive cells and scL-WGS is prone to false negative detection) (Bakker *et al.*, 2015). However, the observed aneuploidy and heterogeneity scores were higher in senescent cells versus young fibroblasts for both FISH and scL-WGS inputs (Supplementary Figure 4), highlighting a trend toward increased numerical chromosomal variability in senescent cells that was present across both methods. These results imply that the ability to detect W-CIN is method-specific and highlight the utility of testing for the presence of W-CIN using an integrated platform that processes multiple experimental inputs in parallel.

## 4 Conclusion

Aneuvis is a web-based application developed to automatically summarize numerical chromosomal variation in single cells. We demonstrate the utility of aneuvis by analyzing the chromosomal status of young and senescent fibroblasts obtained using two techniques: four-colon interphase FISH and scL-WGS. The results from aneuvis show that the differences in W-CIN between treatment groups depend on the experimental method used, and that an integrated framework can highlight trends between experimental methods performed on the same types of cells. Aneuvis allows users to test hypotheses directly within the application and is the first application to summarize numerical chromosomal variation using a graphical user interface. Aneuvis is a user-friendly, web-based, and open-source tool that will enable researchers to identify novel mechanisms underlying the generation of W-CIN.

## Supplementary Information

An online-only supplementary file is available.

